# MOLECULAR STRATIFICATION OF INTESTINAL AND DIFFUSE SUBTYPES OF GASTRIC ADENOCARCINOMA AND ITS PROGNOSTIC IMPACT

**DOI:** 10.1101/2025.09.14.676054

**Authors:** Ronald Matheus da Silva Mourão, Jessica Manoelli Costa da Silva, Valeria Cristiane Santos da Silva, Fabiano Cordeiro Moreira, Paulo Pimentel de Assumpção

## Abstract

Gastric adenocarcinoma is routinely classified by Lauren histology into Intestinal and Diffuse types; however, its prognostic use is limited by molecular heterogeneity within each subtype. We tested whether transcriptomic features refine risk beyond histology and whether biomarker signals generalize across populations. The premise is that when underlying programs diverge across tumors and cohorts, a single biomarker is unlikely to be universally informative; accordingly, integrating molecular signatures with histology may improve risk stratification. We analyzed TCGA-STAD and GTEx gene expression data for discovery and validated the findings in ACRG. Cell-program markers guided unsupervised clustering, and a predictive signature was learned using recursive feature elimination with a random forest and cross-validation. Pathway context was assessed with GSEA, and clinical relevance was evaluated within each Lauren subtype. Clustering of canonical gastric and intestinal programs yielded two molecular clusters that did not explain survival. However, Diffuse tumors with stronger intestinal-like programs tended to experience better outcomes. Consistently, the derived signature was associated closely with Lauren categories and defined two molecular groups. Nevertheless, neither histology nor the signature alone stratified overall survival at conventional thresholds. Moreover, gene-level survival analyses indicated subtype specificity rather than universality. In TCGA, 28 signature genes were associated with overall survival (P < 0.05; HR, 0.79-1.63). ROC analyses for Lauren typing were heterogeneous, with *CRLF1* achieving an AUC of 0.92, followed by *CYP1B1* and *MMRN2*. ACRG gene expression revealed an inversion of the TCGA pattern, with prognostic risk shifting toward the Diffuse subtype. Specifically, *CRLF1, ERG*, and *FRZB* shifted from Intestinal risk to Diffuse risk, while *CDX1* shifted from Intestinal to Diffuse protection. Taken together, these findings suggest that the utility of biomarkers in gastric cancer depends on histology and the population context. Consequently, a contextual strategy that integrates molecular signatures with the Lauren classification has the potential to improve risk stratification beyond either source alone.

## INTRODUCTION

Proposed in 1965, Lauren’s histological classification has served as a key framework for over fifty years in characterizing gastric adenocarcinoma (GAC), offering insights into tumor biology that complement TNM staging [1]. Its categorization into intestinal, diffuse, and mixed subtypes is widely accepted. However, despite its widespread use, there are limitations to the consistent application of this system that impact risk stratification (Kaaijet al., 2020). The main issues include vague definitions of subtype purity, hidden molecular features, and inter-observer variability [3,4]. Without standardized criteria for classifying heterogeneous tumors, different prognostic profiles may be grouped under the same “pure” label (intestinal or diffuse), which reduces the model’s ability to stratify individual patient risk accurately.

The genomic era has begun to explain the molecular basis for Lauren’s histological observations. Robust molecular classifications, such as those proposed by The Cancer Genome Atlas (TCGA) and the Asian Cancer Research Group (ACRG), have demonstrated that Lauren subtypes can comprise biologically distinct entities [5,6]. For example, the TCGA Chromosomal Instability (CIN) subtype is primarily found in intestinal-type tumors, whereas the Genomically Stable (GS) subtype is strongly associated with diffuse-type tumors. Later studies have also shown that tumors with similar histology may exhibit distinct molecular profiles, emphasizing that morphology alone is not enough to understand a patient’s true risk [7].

However, this molecular revolution has not yet been fully applied in clinical practice [8]. High costs, the need for bioinformatics resources, and logistical challenges associated with large-scale sequencing still make it difficult for non-advanced research centers, creating a gap between biological knowledge and practical clinical use [9].

This study aims to bridge this gap by proposing an enhancement to the widely used Lauren system. Our investigation into the prognostic impact of hidden molecular heterogeneity in tumors classified as intestinal or diffuse by standard histology, using gene expression data from reference cohorts, offers a promising approach to stratify high-risk patients within the two main Lauren subtypes.

## MATERIALS AND METHODS

### 1.1 Study cohorts and data acquisition

This analysis used transcriptome data from two independent cohorts. The discovery cohort included public RNA-seq data from gastric adenocarcinoma samples in the TCGA-STAD and normal gastric tissue from the GTEx. Read-count data (STAR-Counts workflow, aligned to GRCh38/hg38) and clinical metadata were obtained via TCGAbiolinks. For independent validation of prognostic findings, we used the ACRG gastric adenocarcinoma cohort with microarray expression data (GEO: GSE62254; platform GPL570), and clinical variables were taken from the original publication.

### 1.2 RNA-seq processing, normalization, and batch correction

The combined dataset (TCGA and GTEx) was filtered to retain only protein-coding genes with detectable expression (i.e., more than 10 total counts). From there, the analysis split into two parts. For differential expression, we used the raw count matrix, with normalization handled internally by the DESeq2 statistical model. For all other analyses (including visualization, clustering, and machine learning), we created a variance-stabilizing transformation (VST) expression matrix, where batch effects between cohorts were corrected using the empirical Bayes method in the limma package.

### 1.3 Differential expression and pathway enrichment

Differential expression analysis employed the Wald test in DESeq2 for the following contrasts: intestinal vs. normal, diffuse vs. normal, intestinal vs. diffuse, and between data-driven groups or clusters. These results informed Gene Set Enrichment Analysis (GSEA) using the clusterProfiler package. Gene lists, pre-ranked by log2 fold change, were evaluated against Reactome, Hallmarks, WikiPathways, and Cell Types (MSigDB v2025.1). Redundancy across enriched terms was reduced using a Jaccard similarity threshold of 0.4.

### 1.4 Development and validation of a molecular gene signature

Molecular subgroups were initially explored through unsupervised hierarchical clustering (Euclidean distance, Ward.D2 linkage) of the VST matrix (z-scores), utilizing a panel of marker genes for gastric and intestinal cell phenotypes (PanglaoDB). We then derived a predictive gene signature for Lauren subtypes via machine learning. Starting from genes differentially expressed between intestinal and diffuse tumors (padj < 0.05, |log2FoldChange| ≥ 1), we applied recursive feature elimination (RFE) with a random forest classifier and 10-fold cross-validation (caret). The dataset was split a priori into 70% training and 30% testing. The final signature was selected using the one-standard-error rule to favor the most parsimonious, robust model.

### 1.5 Prognostic and survival analyses

Survival probabilities were estimated using the Kaplan-Meier method, and the log-rank test was used to assess differences between groups. To stratify patients based on a gene’s continuous expression, the optimal cut-point was determined by maximizing the log-rank statistic (survminer). Associations with survival were quantified as hazard ratios (HRs) from Cox proportional hazards models in univariable and multivariable settings (survival). The classification performance of individual genes was evaluated using receiver operating characteristic (ROC) analysis and the area under the curve (AUC) (pROC).

### 1.6 Statistical analysis

All analyses were performed in R. A P value or FDR ≤ 0.05 was considered statistically significant. Between-group comparisons used the Wilcoxon or Kruskal-Wallis tests as appropriate. Visualizations were generated with ggplot2, pheatmap, and cowplot.

## RESULTS

### Transcriptomic and functional profiles may distinguish gastric cancer subtypes

Beyond morphology, Lauren’s classification exhibits alterations at the molecular level. However, the extent to which this molecular heterogeneity differentiates prognosis in gastric adenocarcinoma remains unclear. To address this, we utilized the TCGA and GTEx datasets (Fig. 1A) to investigate molecular differences between the two classical subtypes.

**Figure 1.**
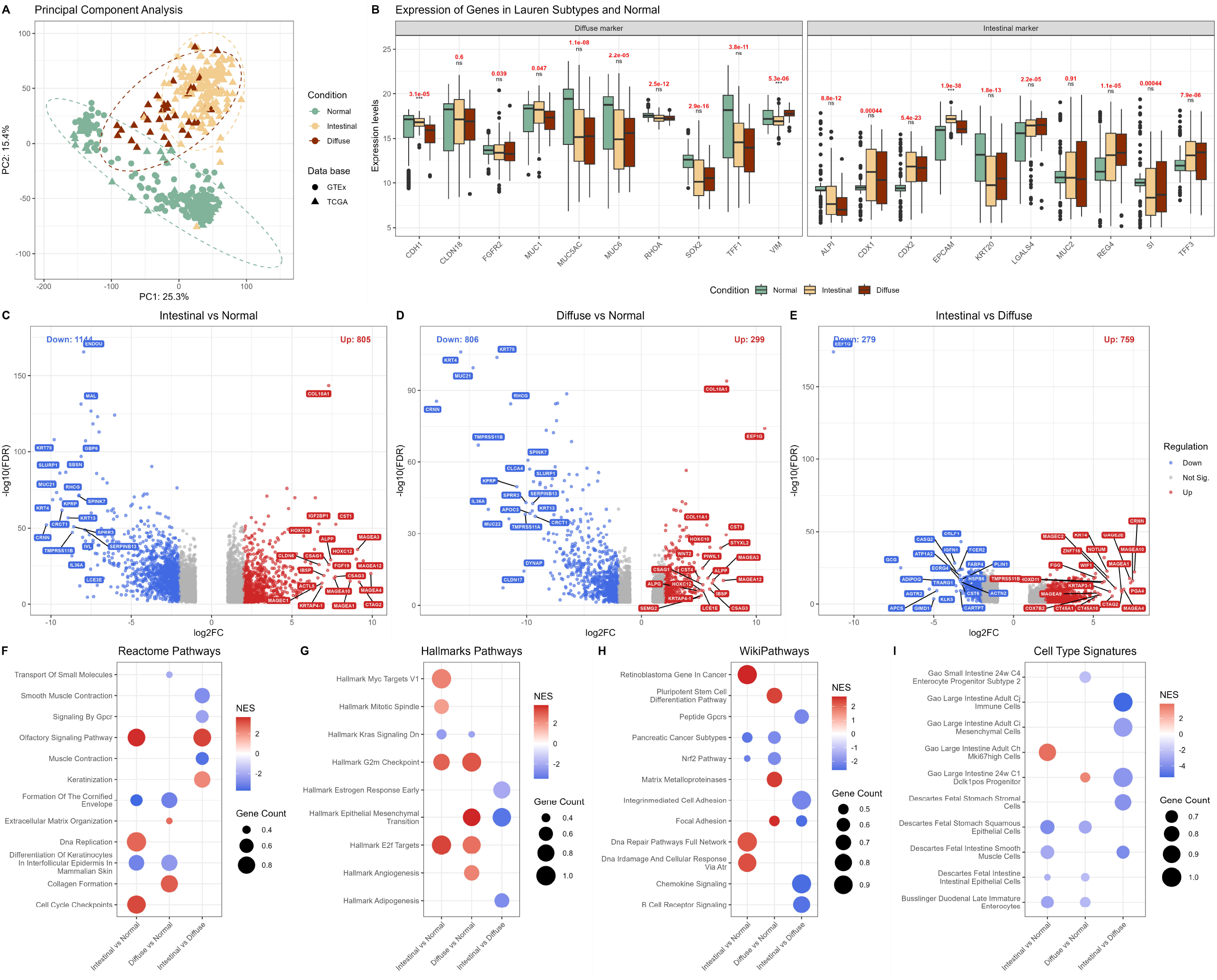
Transcriptomic and functional analysis of gastric adenocarcinoma subtypes. (A) PCA of gene-expression profiles from TCGA (triangles) and GTEx (circles), with colors indicating Normal (green), Intestinal (orange), and Diffuse (brown). (B-D) Volcano plots for the contrasts (B) Intestinal versus Normal, (C) Diffuse versus Normal, and (D) Intestinal versus Diffuse. Axes show log2FC and -log10 FDR; points indicate upregulated genes (red) or downregulated genes (blue). (E) Boxplots of classical markers for Diffuse and Intestinal phenotypes. P values represent differences between tumor subtypes (Wilcoxon test). (F-I) GSEA. Point size indicates gene count, and color encodes NES (red for positive, blue for negative) for: (F) Reactome, (G) Hallmarks, (H) WikiPathways, and (I) Cell Types.

We first examined a canonical marker set commonly linked to the diffuse pattern (*CDH1, CLDN18, FGFR2, MUC1, MUC5AC, MUC6, RHOA, SOX2, TFF1* and *VIM*) and the intestinal pattern (*ALPI, CDX1, CDX2, EPCAM, KRT20, LGALS4, MUC2, REG4, SI* and *TTF3*). Despite notable differences compared to normal tissue, the differences between Diffuse and Intestinal were not significant (Fig. 1B), except for higher *CDH1* and *EPCAM* in the Intestinal type and higher *VIM* in the Diffuse type. To identify subtype-specific profiles, we conducted differential expression analysis for Intestinal vs. Normal (Fig. 1C), Diffuse vs. Normal (Fig. 1D), and Intestinal vs. Diffuse (Fig. 1E). The comparison of Intestinal vs. Normal showed the greatest change, with 805 genes upregulated and 1,144 downregulated. The Diffuse vs. Normal comparison revealed a distinct profile, with 299 genes upregulated and 806 downregulated. In the direct comparison of Intestinal vs. Diffuse, 759 genes were more expressed in the Intestinal type, while 279 genes were higher in the Diffuse type.

GSEA showed that, compared to normal tissue, both subtypes are enriched (positive NES) for cell proliferation programs such as Cell Cycle and Cell Cycle Checkpoints (Reactome, panel F) and the Hallmark G2M Checkpoint (panel G). The direct comparison between Intestinal and Diffuse subtypes revealed different patterns: Diffuse is characterized by enrichment in EMT (Hallmarks, panel G), along with Extracellular Matrix Organization (Reactome, panel F), Intermediate Cell Adhesion (WikiPathways, panel H), and Integrin-mediated Cell Adhesion (WikiPathways, panel H). In contrast, both subtypes were enriched for intestinal cell signatures, including Descartes Fetal Intestinal Epithelial Cells, Gao Large Intestine Adult CI Mesenchymal Cells, and Gao Large Intestine Adult CI Immune Cells (panel I). This molecular heterogeneity, where expression profiles indicate intestinal phenotypes within diffuse tumors, suggests that a gene signature-based reclassification could have clinical implications.

### Development of a molecular signature for prognostic stratification

To address the question raised above, we looked for molecular subgroups with an intestinal-like profile and potential clinical relevance. Using unsupervised hierarchical clustering based on markers of canonical gastric and intestinal cell-expression phenotypes, samples divided into two major molecular groups with no link to Lauren subtype (χ^2^=0.0028, p-value=0.95; n=196; Fig. 2A). Expression programs from the Huang signature (2025), an intestinal gastric-cell signature, along with enterocyte markers, mainly drove this separation, which we call the intestinal signature. Consistently, genes that were differentially expressed between clusters (C2 vs C1) indicated the presence of an intestinal-cell profile in C2 (Fig. 2B). Supporting these results, Lauren type alone did not show prognostic value (p-value=0.23; Fig. 2C), nor did stratifying Lauren types by the molecular clusters identified in Fig. 2A (p-value=0.24; Fig. 2D).

**Figure 2.**
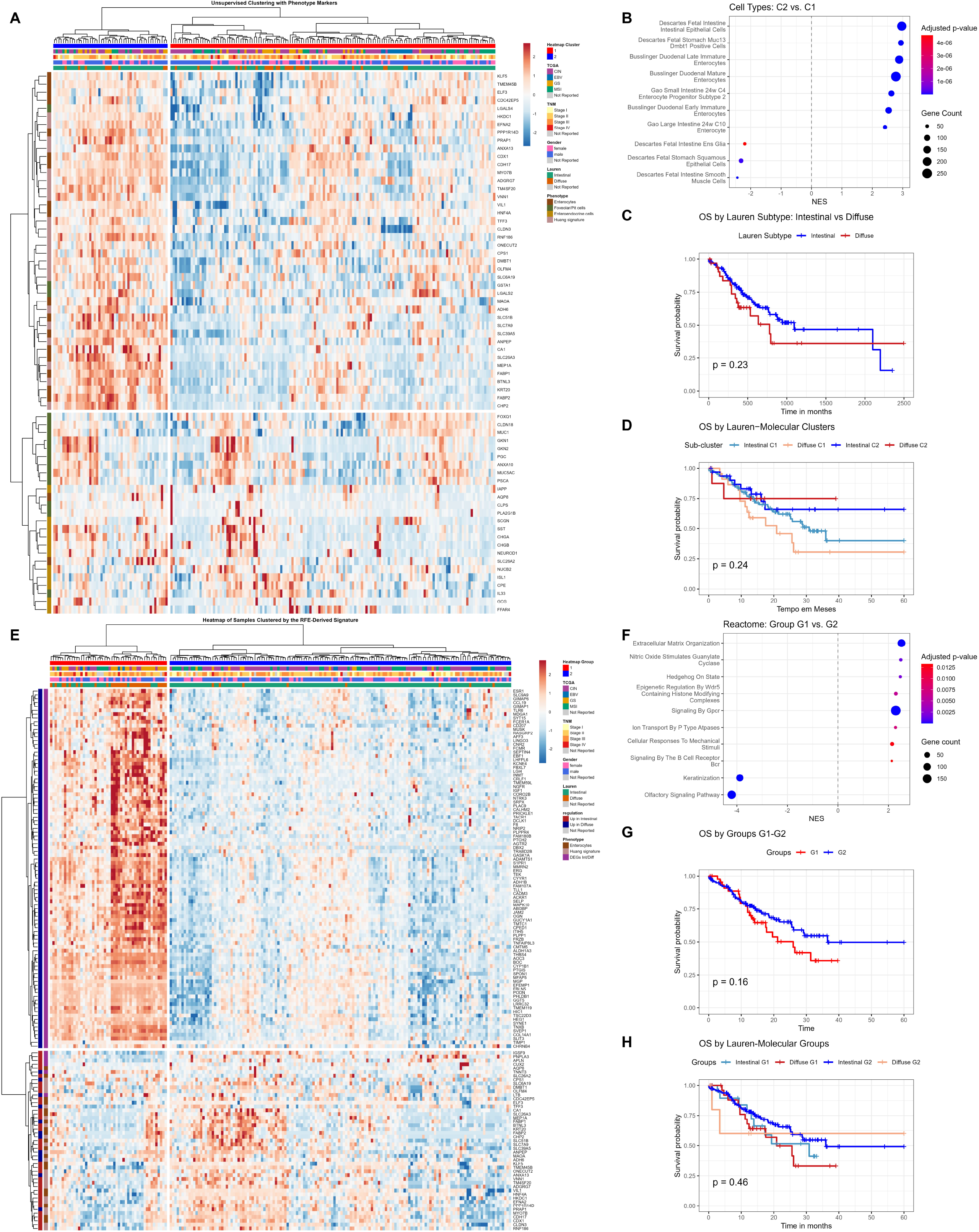
Integrated molecular signatures and prognostic relevance in gastric adenocarcinoma (A) Hierarchical-clustering heatmap of tumor samples based on a panel of marker genes for cellular phenotypes. Top annotations indicate the Lauren classification and clinical features. Matrix colors represent gene-expression z-scores, where red indicates high values and blue indicates low values. (B) Pathway enrichment (GSEA) between the molecular clusters identified in (A). (C) Kaplan-Meier curve for overall survival. (D) Kaplan-Meier curve stratified by clinico-molecular subgroups derived from integrating the clusters in (A) with the Lauren classification. (E) Heatmap guided by a machine-learning-refined gene signature (RFE) that segments samples into three new molecular clusters. (F) GSEA comparing the clusters with the highest and lowest expression of the signature identified in (E). (G) Kaplan-Meier curve stratified by the three molecular clusters in panel (E). (H) Final Kaplan-Meier curve stratified by four subgroups integrating the Lauren classification with High/Low expression of the refined gene signature.

Although stratification by molecular profile was not statistically significant, combining the molecular clusters with the Lauren classification revealed distinct prognostic patterns (Fig. 2D). The group of patients with histologically Diffuse tumors in cluster C2, characterized by high expression of the intestinal signature (Huang signature plus Enterocytes), exhibited the most favorable prognosis overall. Conversely, patients with histologically Diffuse tumors in C1, marked by low intestinal-phenotype expression, had the lowest survival rates compared to all other groups. Overall, these findings suggest that molecular diversity within Lauren subtypes may have clinical significance, indicating that an intestinal-like profile could mitigate the negative prognosis typically associated with the Diffuse subtype.

Given the limitations of the exploratory approach described earlier, we next derived a gene signature from the differentially expressed genes between Intestinal and Diffuse tumors, using RFE with a random forest to select the most predictive features. Clustering samples based on this new signature resulted in a more coherent classification into two molecular groups and showed a clear association with Lauren subtypes (χ^2^=63.70, p-value = 1.448e-15; n=196; Fig. 2E). Functional differences were confirmed by GSEA (Fig. 2F), which compared the higher-expression group (G1) with the lower-expression group (G2) of signature genes.

Although the two groups alone were not linked to survival (p-value = 0.16; Fig. 2G), combining this signature with the Lauren classification showed promising prognostic patterns. The final division into four clinico-molecular groups, considering Groups 1 and 2, revealed a trend toward separation of survival curves that did not reach statistical significance (p-value = 0.14; Fig. 2H). Patients with either Diffuse or Intestinal type tumors and high signature expression (G1) had the lowest survival probabilities compared to those in the Intestinal G2 group. Conversely, patients with intestinal tumors and low expression of the diffuse-tumor differential signature (G2) experienced the best outcomes. This finding suggests that the clinical importance of the signature is unlikely to be universal across all genes, prompting further investigation into the strength of the association between overall survival and the expression of each gene highlighted in panels 2A and 2F.

### Identification and validation of prognostic biomarkers with subtype-specific value

To identify the specific markers contributing to this heterogeneity, we conducted gene-level survival analysis. In the TCGA cohort, univariable Cox regression identified 28 signature genes linked to overall survival (P<0.05), with HR values ranging from 0.79 to 1.63 (Fig. 3A). Among these, 25 functioned as higher-risk markers (HR>1), and 3 appeared protective (HR<1). Considering the effect size, small effects were most common (HR 1.05 to 1.20 / 0.95 to 0.83), followed by moderate effects (HR 1.20 to 1.50 / 0.83 to 0.67). *CMTM5* significantly impacted risk direction (HR ≥ 1.50 / ≤ 0.67). On the protective side, *KLF5* and *TMEM45B* showed moderate associations, while *CDX1* exhibited a small effect. The genes displayed in Fig. 3A are selected as promising molecular markers.

**Figure 3.**
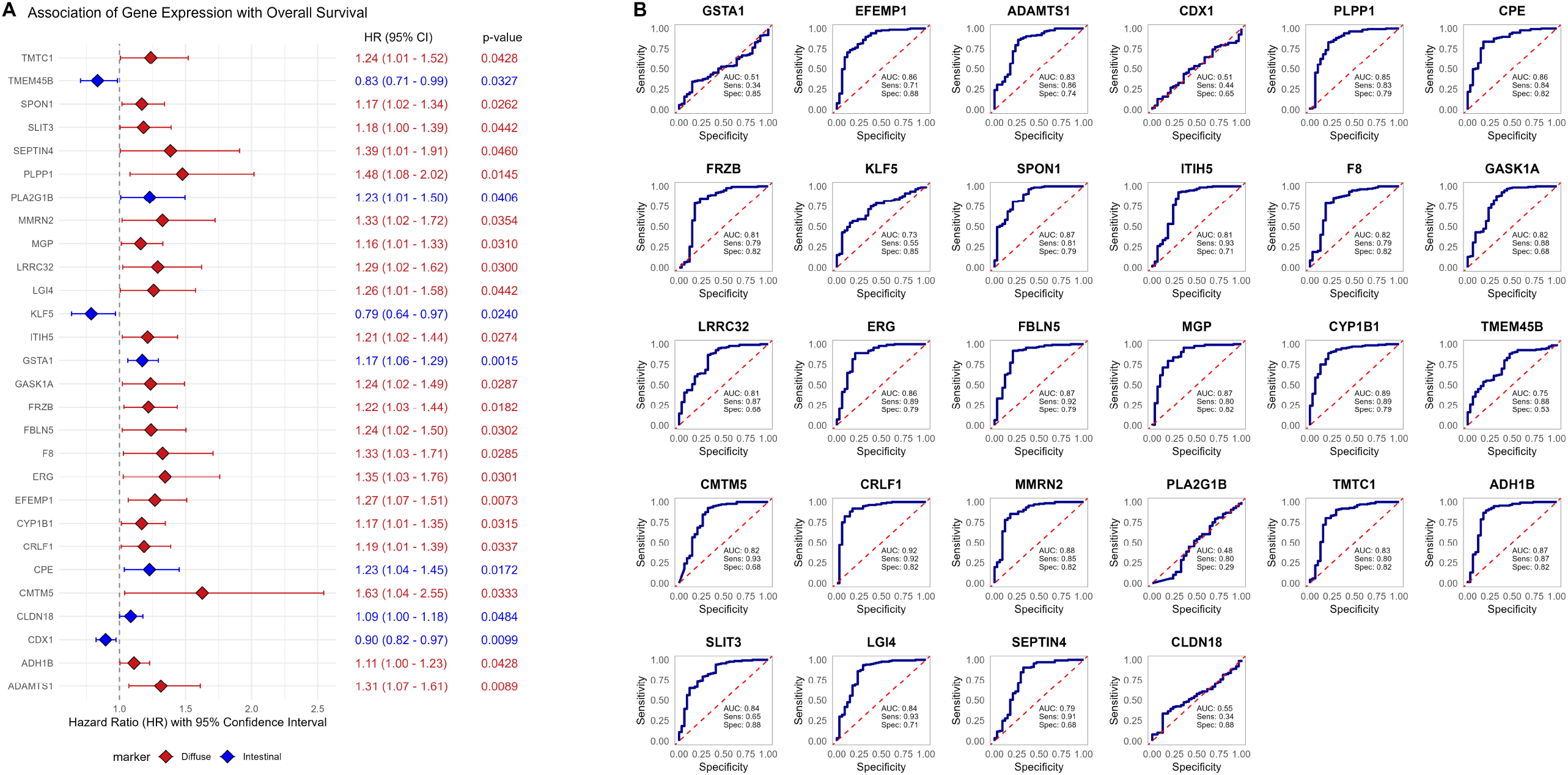
Identification of a subtype-specific prognostic signature. TCGA cohort: (A) Forest plot of univariable Cox analysis for overall survival. (B) ROC curves evaluating the predictive ability of individual genes for Lauren subtypes.

Meanwhile, ROC curves differentiating the two canonical Lauren types exhibited varied performance across these potential markers (Fig. 3B). *CRLF1* was the best classifier (AUC = 0.92), followed by CYP1B1 (AUC = 0.89) and *MMRN2* (AUC = 0.88). An additional 18 genes had AUCs between 0.81 and 0.87, including *SPON1, FBLN5, MGP, ADH1B, EFEMP1, CPE, ERG, PLPP1, SLIT3, LGI4, ADAMTS1, TMTC1, F8, GASK1A, CMTM5, FRZB, ITIH5*, and *LRRC32. SEPTIN4, TMEM45B*, and *KLF5* scored below 0.80, while *CLDN18, GSTA1, CDX1*, and *PLA2G1B* showed AUCs of 0.55 or less.

To assess the prognostic value of these genetic markers within each Lauren subtype, we conducted Cox survival analyses separately for Intestinal and Diffuse tumors. In the TCGA cohort, the univariable results showed that prognostic associations were mainly clustered in the Intestinal subtype (Fig. 4A). Among the genes with significant associations, 75.0% acted as risk factors and 7.1% as protective factors—all exclusively observed within this subtype, with no relevant associations found in the Diffuse subtype. Notable risk markers in the Intestinal subtype included *ADAMTS1, EFEMP1, SLIT3, CPE, LGI4, TMTC1, GASK1A*, and *FRZB*, while *KLF5* and *CDX1* served as protective factors. F8 was the only gene identified as a risk factor in the Diffuse subtype. The corresponding heatmap (Fig. 4B) clearly shows a grouping of Diffuse tumors, with most genes exhibiting higher expression. This pattern, except for genes like *CDX1, KLF5*, and *CLDN18* (which is more highly expressed in the intestine), underscores that prognostic interpretation must be specific to each subtype, as it does not directly correspond to expression levels.

**Figure 4.**
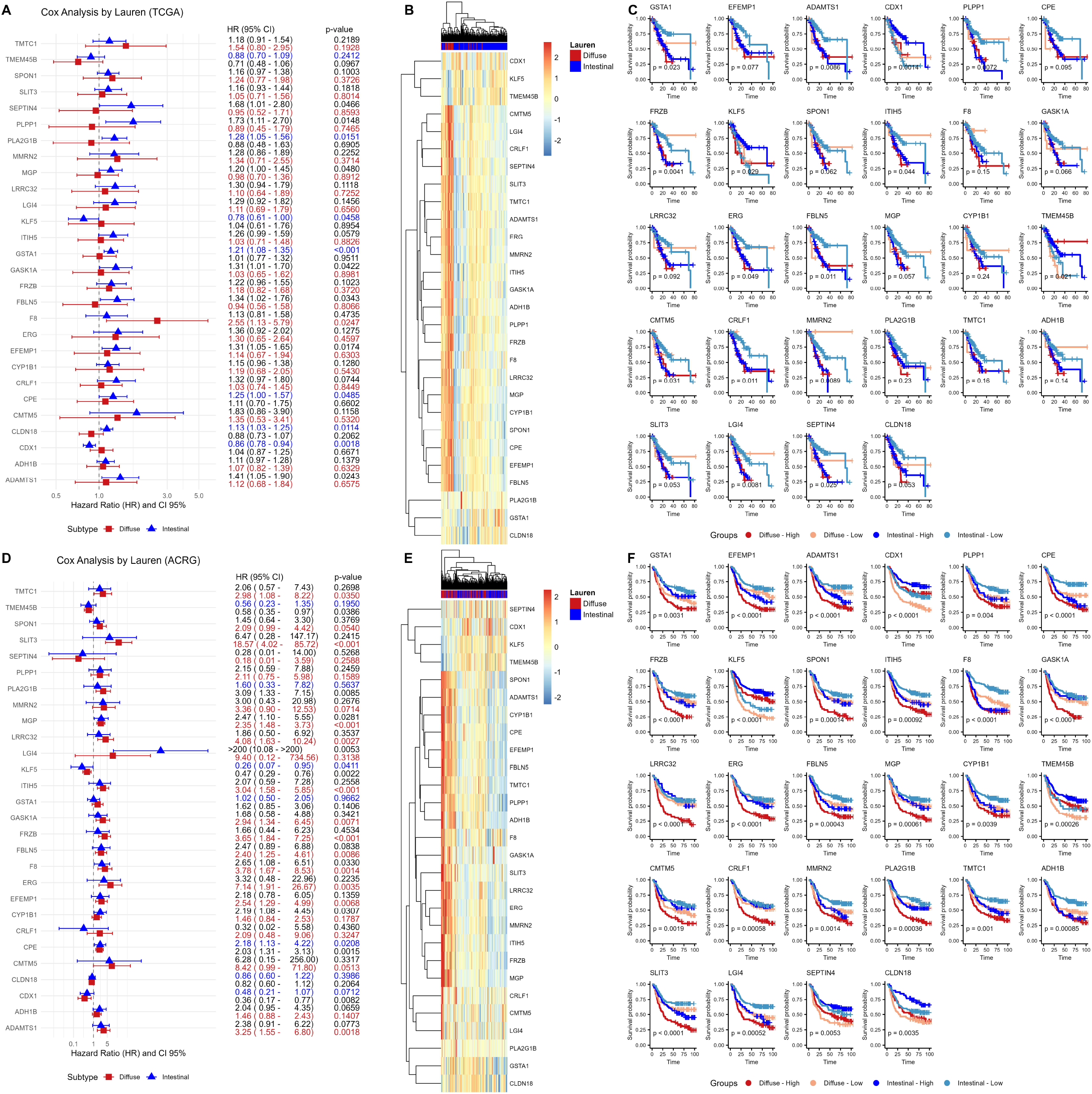
Subtype-specific survival analysis for the Lauren classification and external validation. (A) Univariable Cox regression in the TCGA cohort showing HRs for the Intestinal and Diffuse subtypes. The prognostic value is concentrated in the Intestinal subtype. (B) Heatmap of the expression of the 28 genetic markers in TCGA, with samples grouped by Lauren subtype. (C) Kaplan-Meier curves for representative genes in TCGA. Patients are stratified by subtype (Intestinal, Diffuse) and expression level (High, Low). (D) Cox regression in the ACRG validation cohort, indicating a shift in the prognostic profile toward the Diffuse subtype. (E) Gene-expression heatmap in ACRG. (F) Kaplan-Meier curves for representative genes in ACRG, demonstrating subtype-specific survival impact in this cohort. In (C) and (F), P values refer to the log-rank test comparing the four groups.

In the TCGA cohort, Kaplan-Meier curves supported these findings (Fig. 4C). Patients with Intestinal tumors and high expression of risk genes (Intestinal-High) had significantly worse survival than those with low expression (Intestinal-Low). Consistently, Intestinal-Low showed the best prognosis, often not inferior to that of the Diffuse groups, regardless of gene expression level. For protective markers in the Intestinal subtype (*KLF5* and *CDX1*), the pattern reversed, with Intestinal-High outperforming Intestinal-Low, while Diffuse groups showed intermediate survival or no clear difference. For example, *SEPTIN4* had a p-value of 0.025 in the overall log-rank test and an HR greater than 1.5 in the Intestinal subtype, confirming poorer outcomes in the Intestinal-High compared to the Intestinal-Low subtype.

When we replicated the analysis in the non-Western ACRG cohort (Fig. 4D), we observed a notable inversion in association patterns, with prognostic risk shifting toward the Diffuse subtype. In this cohort, 39.3% of significant genes indicated risk and 10.7% indicated protection in Diffuse, while 28.6% indicated risk and 3.6% indicated protection in Intestinal. Compared to TCGA, the changes followed three main trends. First, some genes expanded their risk profile, as shown by *ADAMTS1, EFEMP1*, and *SLIT3*, which shifted from being associated with risk in only the Intestinal subtype to being associated with risk in both subtypes. Second, there was subtype migration with the effect direction remaining consistent; for example, *CRLF1, ERG*, and *FRZB* moved from a risk association in the Intestinal subtype (TCGA) to a risk association in the Diffuse subtype (ACRG), and *CDX1* moved from a protective association in the Intestinal subtype to a protective association in the Diffuse subtype. Third, new genes became significantly linked to survival, such as *ADH1B* and *F8*, which confer risk in the Intestinal subtype, and *PLA2G1B*, which confers risk in the Diffuse subtype. The ACRG heatmap (Fig. 4E) also showed samples clustering by Lauren subtype, with higher expression levels predominating in Diffuse for most genes.

Kaplan-Meier analyses in ACRG confirmed this new subtype-specific pattern (Fig. 4F). Patients with Diffuse tumors and high expression of the corresponding risk genes (Diffuse-High) had the poorest survival. In contrast, the Diffuse-Low group showed better prognosis within the subtype and often outperformed the Intestinal groups. For protective genes in Diffuse (*KLF5, TMEM45B*, and *CDX1*), Diffuse-High demonstrated better survival than Diffuse-Low, which still exceeded the outcomes of the Intestinal groups. For risk genes in the Intestinal subtype (*CPE* l and *ADH1B*), the Intestinal-High group again had the worst outcomes compared with the Intestinal-Low group, ranking among the lowest across the four groups.

## DISCUSSION

Lauren’s histological classification remains a cornerstone for characterizing gastric adenocarcinoma, yet its prognostic utility is consistently undermined by substantial molecular heterogeneity within each subtype [1,5,6]. Our study directly addresses this gap, demonstrating that the predictive value of genetic markers associated with Lauren categories is not universal but strongly conditioned by both histological context and population background. This finding challenges the feasibility of a uniform, one-size-fits-all biomarker for gastric cancer. It reinforces the emerging consensus that classification systems must move beyond morphology to integrate molecular information.

Our transcriptomic analysis showed that, although Intestinal and Diffuse tumors display distinct global expression patterns, Diffuse tumors frequently harbor intestinal-like gene programs. An exploratory clustering analysis based on canonical gastric and intestinal cell markers was unable to stratify survival. However, diffuse tumors enriched with intestinal signatures tended to exhibit improved outcomes. This observation parallels recent integrative frameworks that identified overlap between histology and molecular subtypes: Intestinal tumors frequently align with CIN or MSI subtypes, while Diffuse tumors are enriched in the GS or MSS/EMT subtypes, the latter characterized by epithelial-mesenchymal transition and poor prognosis [5,6].

Recent studies confirm that integrating Lauren with TCGA or ACRG subtypes yields clinically meaningful refinements. For instance, Biesma et al. (2024) demonstrated that CIN-diffuse tumors had a five-year survival rate of only 31%. In contrast, GS-intestinal tumors had a rate of 61%, establishing that the intersection of morphology and molecular status can define highly divergent outcomes [10]. Our results are concordant: neither Lauren’s gene signature nor our machine-learning gene signature alone stratified survival with statistical significance; yet, their integration improved the delineation of risk groups, underscoring the added value of combining histological and molecular frameworks.

At the gene level, prognostic effects were strikingly subtype-specific. In TCGA, 25 of 28 significant genes correlated with survival only in intestinal tumors. For example, *ADAMTS1* acted as a risk gene exclusively in this subtype, whereas *KLF5* and *CDX1* were protective, despite similar expression in diffuse tumors. This suggests that the prognostic effect of a gene depends on the biological circuitry of its host context. Comparable findings were reported by Bao et al., who noted that Lauren types alone were non-prognostic and identified high-risk diffuse tumors [11]. In our analysis, *CRLF1* achieved an AUC of 0.92 for distinguishing Lauren types, further highlighting the diagnostic but not universal prognostic value of individual genes. These results align with broader evidence showing that single-gene biomarkers have limited reproducibility in gastric cancer, whereas multi-gene panels capture the complexity of tumor biology and yield stronger predictive power [12].

Importantly, validation in the ACRG cohort revealed a profound inversion of the TCGA pattern: many markers that predicted outcome in Intestinal tumors in TCGA shifted prognostic impact to the Diffuse subtype in *ACRG. CRLF1, ERG*, and *FRZB* migrated from Intestinal risk genes to Diffuse risk genes, while *CDX1* shifted from Intestinal to Diffuse protection. Such reversals mirror other reports of geographic and population-specific variability in gastric cancer. Large comparative studies have shown that molecular subtype distributions differ between Eastern and Western patients, with Western cohorts showing predominance of CIN and lower MSI frequency in early stage and similarities in late stage [13].

Lauren classification has maintained its independent prognostic value in recent Asian cohorts, where the intestinal subtype is linked to better outcomes, as demonstrated in a Chinese study using a modified Lauren system [14] and in immune-profiling work [15] showing poorer survival in diffuse-type versus intestinal-type tumors. However, its prognostic power is less consistent when used without molecular stratification, as shown in the study by Biesma et al. (2024). Combining Lauren with TCGA GS/CIN molecular subtypes significantly improved risk discrimination compared with Lauren histology alone [10] Collectively, these findings support the conclusion that external factors such as etiology, diet, and H. pylori prevalence, along with intrinsic genetic backgrounds, influence the prognostic impact of biomarkers, as confirmed in the two cohorts analyzed in our study.

In summary, our study reinforces the idea that refining Lauren’s classification for prognostic use requires a contextual approach. The value of molecular biomarkers in gastric cancer depends not only on the histological subtype but also on the population being studied. Universal biomarkers have consistently failed to produce reliable results, whereas subtype-specific and cohort-focused panels have shown greater potential. The next step is to validate these subtype-specific panels using clinically feasible assays, such as immunohistochemistry, and to test them in prospective, multicenter cohorts further. This way, the histological framework of Lauren can develop from a descriptive classification into a more actionable clinical tool that guides prognosis and treatment decisions within specific populations.

## ACKNOWLEDGMENTS

We are grateful to CAPES (Coordenação de Aperfeiçoamento de Pessoal de Nível Superior) for granting a doctoral fellowship to the authors. Financial support from Fundação Amazônia de Amparo a Estudos e Pesquisas (FAPESPA) is also appreciated.

## COMPETING INTERESTS

The authors declare that they have no conflicts of interest.

## AUTHOR CONTRIBUTIONS

R.M.S.M. were responsible for designing the statistical analyses, interpreting results, and writing the draft. J.M.C.S., and V.C.S.S. contributed to data and manuscript revision. P.P.A. and F.C.M. supported the conceptual development of the study.

## Notes

### Competing Interest Statement

The authors have declared no competing interest.

